# The dynamic Nexus: Gap junctions control protein localization and mobility in distinct and surprising ways

**DOI:** 10.1101/2020.04.06.027540

**Authors:** Sean McCutcheon, Randy F. Stout, David C. Spray

## Abstract

Gap junction (GJ) channels permit molecules, such as ions, metabolites and second messengers, to transfer between cells. Their function is critical for numerous cellular interactions. GJ channels are composed of Connexin (Cx) hexamers paired across extracellular space and typically form large rafts of clustered channels, called plaques, at cell appositions. Cxs together with molecules that interact with GJ channels make up a supramolecular structure known as the GJ Nexus. While the stability of connexin localization in GJ plaques has been studied, mobility of other Nexus components has yet to be addressed. Colocalization analysis of several nexus components and other membrane proteins reveal that certain molecules are excluded from the GJ plaque (Aquaporin 4, EAAT2b), while others are quite penetrant (lipophilic molecules, Cx30, ZO-1, Occludin). Fluorescence recovery after photobleaching (FRAP) of tagged Nexus-associated proteins showed that mobility in plaque domains is affected by mobility of the Cx proteins. These novel findings indicate that the GJ Nexus is a dynamic membrane organelle, with cytoplasmic and membrane-embedded proteins binding and diffusing according to distinct parameters.

**Summary Statement:** Gap junctions are clustered membrane channels in plasma membrane of astrocytes and other cells. We report new information on how gap junctions control location and mobility of other astrocyte proteins.

## Introduction

Gap junction (GJ) channels are paired hexameric structures composed of connexin (Cx) proteins^1^. More than twenty Cx isoforms are expressed in specific patterns in most cell populations throughout the body. Gap junction plaques are structures made up of aggregated gap junction channels, and interactions between connexins and other integral membrane and non-membrane tethered proteins make up the gap junction Nexus^2^. The gap junction plaque has been regarded as a highly organized, rigid contact area between cells, and its biochemical integrity and close particle packing enabled the pioneering isolation, purification and x-ray diffraction studies of its structure ^3^. Recent live cell imaging of fluorescent protein tagged connexins has revealed that while some GJ plaques are quite stable over minutes of observation, including those comprised of the most commonly expressed gap junction protein Cx43, others such as Cx30, Cx26, and Cx36 are more fluid and can rearrange within tens of seconds^4,5^. Of note, Cx43 GJ plaque stability is governed by cysteine interactions at the cytoplasmic carboxyl-terminus as evidenced by increased fluidity when Cx43 was truncated at amino acid 258, and after replacement of cytoplasmic cysteine residues with alanine. Moreover, Cx43 mobility was found to be rapidly modifiable by cell-permeable reducing agents, indicating a dynamic and reversible process^6^.

It has long been recognized that gap junctions exist as plaques, and more recently it has been appreciated that other proteins bind to connexins to create a Nexus of molecules critical to intercellular signaling. However, the degree to which these interactions are stable and dictate the distribution of other channels, membrane components, and scaffolding molecules has not been thoroughly evaluated. Membrane-embedded proteins /molecules are constrained in their movement by the presence and organization of other proteins that interact within the membrane and by proteins that form cytoplasmic scaffolding or extracellular aggregating lattices. As shown by freeze fracture electron microscopy, core transmembrane proteins of tight and gap junctions are often intermixed, with enastomosing tight junction (TJ) strands encompassing gap junction plaques. Direct interaction between core proteins of each junction type have been reported^7,8^, and binding of Cx43 to PDZ domain of the TJ adaptor protein zonula occludens-1 (ZO-1) is believed to play a role in regulating GJ size^9^. In addition, interdigitation of gap junction and tight junction particles at the margin between apical and basolateral domains of polarized cells creates a potential signaling domain where intercellular signaling molecules might regulate extracellular tightness. However, impact of GJ-TJ molecular interactions on overall GJ plaque structure, stability and dynamics has not been fully elucidated.

To gain insight into the dynamic relationship between Cx43 and other cellular components, we have determined the diffusivity and location of other molecules relative to the Cx43 plaque. For these studies we selected for comparison small fluorescent lipids and lipid-tethered fluorescent proteins, the water channel Aquaporin4 (AQP4) whose presence with Cx43 at astrocyte endfeet regulates ion homeostasis, a glutamate transporter (EAAT2b) that provides astrocytes with a mechanism to take up glutamate at active synapses, and junction-associated proteins (the other astrocyte gap junction protein Cx30, the tight junction proteins occludin and ZO-1).

Fluidity of gap junction channels within the plaque and stability of interactions between connexins and a few of their binding partners have been quantified in previous studies by fluorescence recovery after photobleaching (FRAP)^10^. Here we examine the mobility of a variety of membrane-embedded proteins and other gap junction Nexus components in the presence of stable and fluidized Cx43 gap junctions. We tested whether Cx43 plaques exclude some molecules and permit diffusion of others. We tested if non-Cx43 molecules that enter the GJ plaque area are mobile. Finally, we tested whether increasing fluidity of the Cx43 plaques by C-terminus truncation or altering cytoplasmic cysteine residues (C260, 271, 298 mutated to alanine) increased fluidity of the GJ-associated proteins.

## Results

### Gap junction plaques modify localization of small membrane-associated molecules

The localization of membrane-associated molecules with respect to Cx43 GJ plaques was examined by two-channel confocal microscopy (see Fig 1A-C) and degree of overlap was quantified with Pearson correlation coefficients obtained from line scans along the junctional membrane (Fig 1 D-E). The coumarin-linked lipid CC2-DMPE was found to be distributed throughout the cells, labeling both plasma and intracellular membranes (Fig. 1A). It was also distributed rather uniformly in plaque and nonplaque domains, as determined by line-scans of the appositional membranes at the margin between plaque and nonplaque as identified with fluorescence of msfGFP-Cx43 (rat Cx43 with a monomerized GFP).

**Figure 1.**
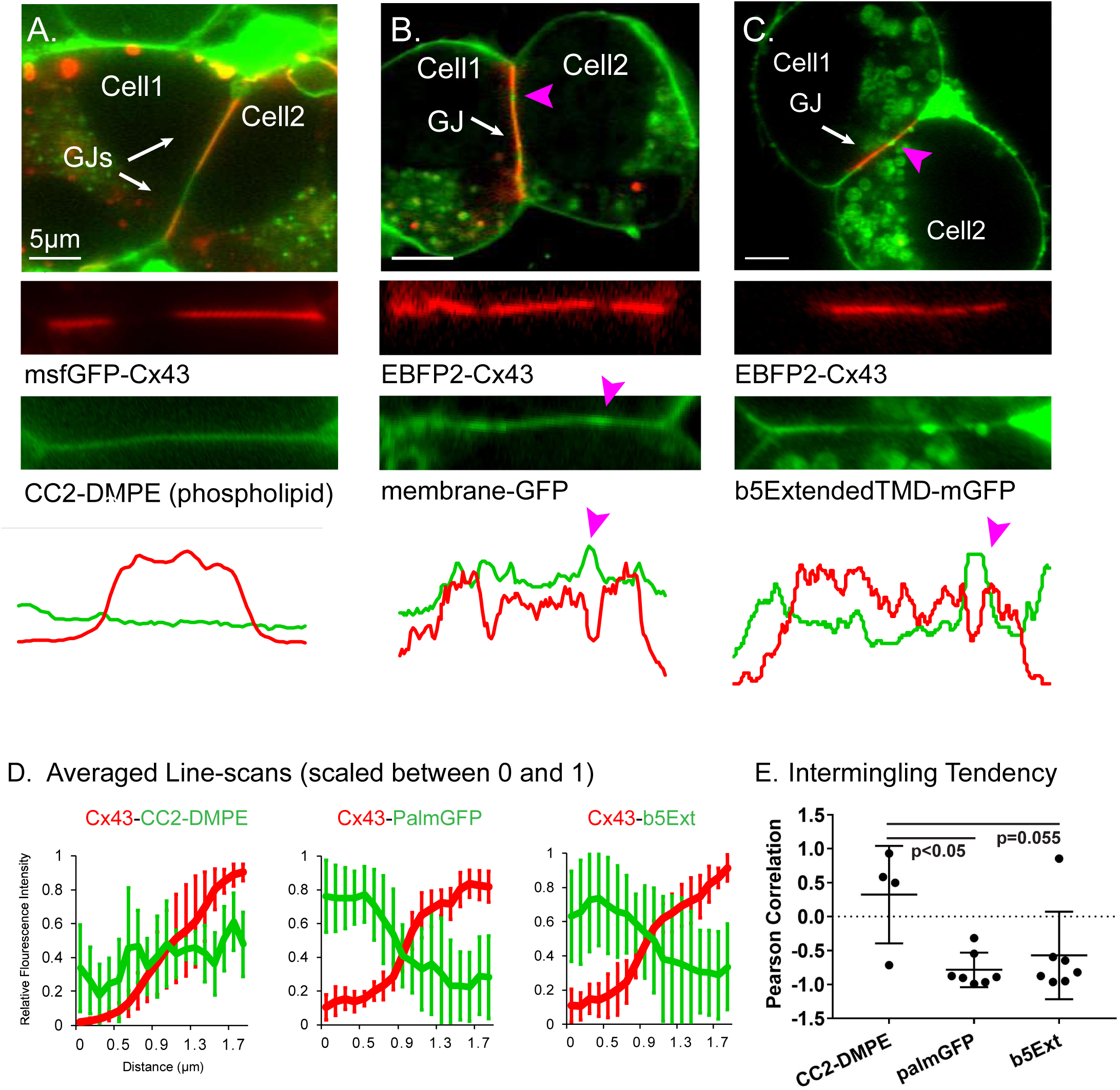
Gap junction plaques modify localization of membrane-bound molecules. **A)** Fluorescence micrographs of N2A cells co-expressing fluorescently-tagged small membrane molecules with fluorophore tagged-Cx43, illustrating that phospholipid CC2-DMPE penetrates Cx43 plaques (white arrows) and has an unaltered distribution vs the non-plaque region. Individual channels are shown below the overlay images. For panels in “A” msfGFP-Cx43 is pseudocolored red for consistency with EBFP2-Cx43 which is red pseudocolored throughout the following figures. Line-scans spanning the entire plaque for the two channels are shown below the example images. **B)** membrane-tethered GFP and **C)** transmembrane protein b5Ext penetrate the plaque but have a lower density than in non-plaque regions. **D)** Average line scans of relative fluorescence intensity for fluorophore-tagged molecules at the edge of Cx43 plaques. n ≥ 4 **E)** Pearson correlation, or tendency to intermingle with Cx43 at the plaque. Magenta arrows in B, C indicate void spaces in the Cx43 plaque.

By contrast to the rather uniform distribution of the purely lipid probe throughout the cell membrane, the two small labeled membrane-embedded proteins that we examined (membrane localized, palmitoylated GFP and a single-pass transmembrane protein b5Extended-GFP) showed uniform distribution within the non-junctional plasma membrane but appeared to be partially excluded from the plaque. For these experiments EBFP2-tagged rat Cx43 was expressed to allow visualization of the gap junction plaque. Distribution of membrane-tethered GFP (Figure 1B1) and b5Ext-GFP (Figure 1C1) was variable within the plaque region. We observed higher localization of both membrane-bound GFP proteins to areas where holes were present in the gap junction plaque in both images and line scans as indicated by magenta arrowheads in Figure 1. Averaged arbitrary intensity values for phospholipid and membrane-tethered components are shown for line scans across the transition from non-plaque membrane to plaque membrane (Figure 1D). Intermingling tendency of the phospholipid and membrane tethered fluorescent-tagged molecules within the gap junction plaque was assessed as the Pearson correlation of fluorescence across the two color-channels. This analysis indicated that presence within the plaque was slightly higher than in nonjunctional membrane for CC2-DMPE (0.32±0.62), whereas membrane tethered GFP and b5Ext were largely excluded from the plaque (r=-0.78±0.23 and -0.57±0.60, respectively) (Figure 1E).

We also examined the distribution of several multi-pass membrane-embedded proteins compared with that of Cx43 in plaques composed of EBFP2-Cx43. For this, we visualized Cx43 simultaneously with the water channel AQP4, the glutamate transporter EAAT2b, the other major astrocyte-expressed gap junction protein Cx30, and the tight junction protein occludin (Ocln), as well as the cytoplasmic junction associated protein ZO-1 (Figure 2). AQP4 and EAAT2b showed similar distribution, being largely excluded from the Cx43 GJ plaque. Pearson coefficients for these two proteins were less than -0.5 (−0.66±0.24 for AQP4 and -0.70±0.19 for EAAT2b) as shown in Figure 2C. By contrast, Cx30 and Ocln fluorescence largely overlapped with Cx43 within the plaque. The overlapping distribution of these proteins was shown in line scans where local discontinuities (lacunae in the GJ plaque) were apparent in both traces (Figure 2A, arrows). Pearson coefficients were nearly +1, 0.88±0.09 for Cx30 and 0.77±0.23 for Ocln (Fig 2C, Rightmost graph). Cx43 binds the scaffolding protein ZO-1 through interaction between amino acids at the end of the carboxyl terminus of Cx43 and the PDZ-2 domain of ZO-1 ^11,12^. A fluorescent tag appended to the carboxyl terminus of Cx43 disrupts the binding site for ZO-1^13^; thus we again used our EBFP2-Cx43 construct with its free carboxyl-terminus to visualize the localization of Cx43 and ZO-1 in live cells. ZO-1 showed a pattern of distribution that overlapped with that of Cx43 (Pearson coefficient = 0.72±0.20).

**Figure 2.**
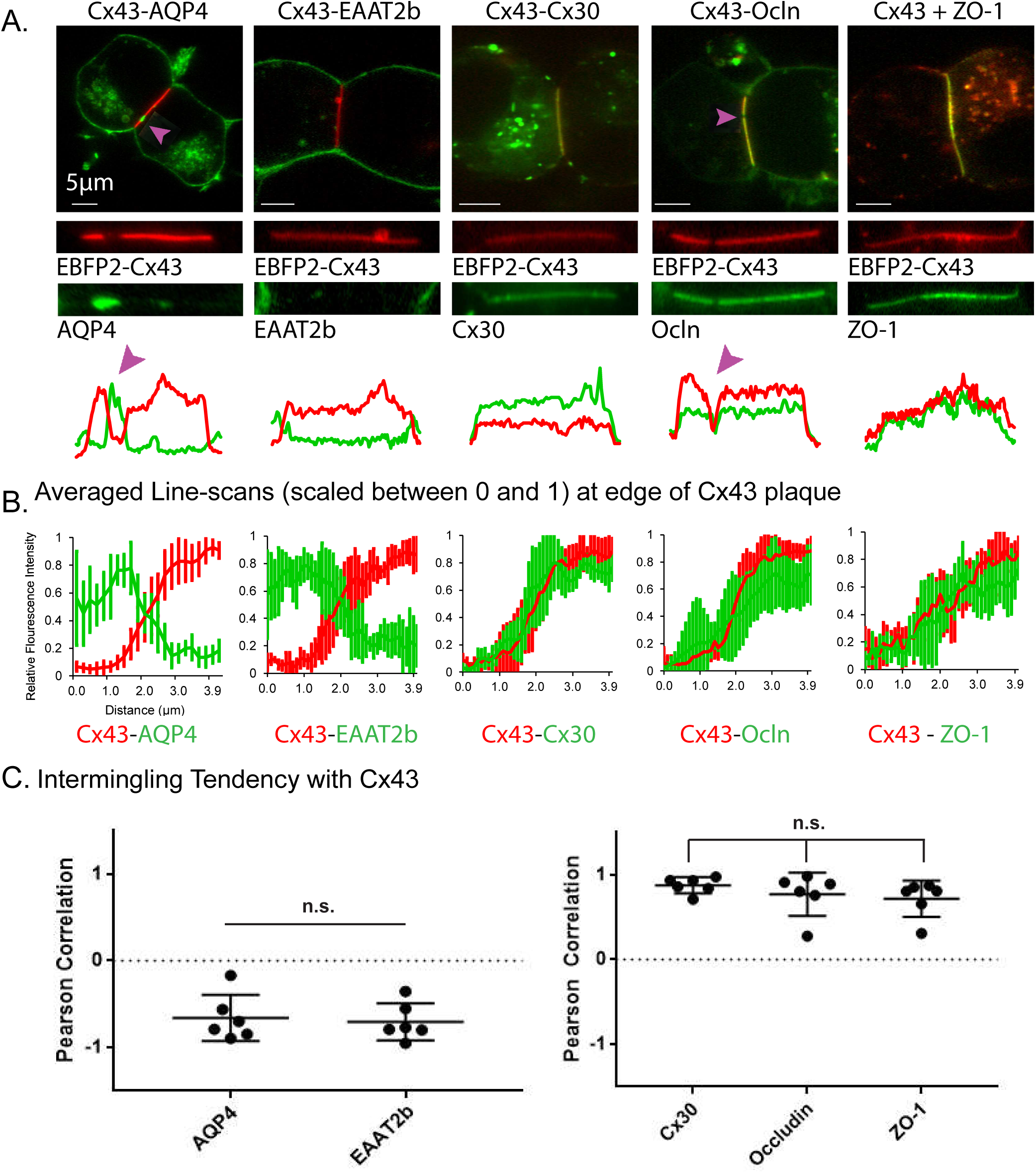
Integration or exclusion of membrane proteins from gap junction plaques. **A)** Fluorescence micrographs of N2A cells co-expressing GFP-tagged membrane proteins and BFP-tagged Cx43 demonstrating that AQP4 and EAAT2b are excluded from Cx43 plaques, whereas Cx30, Occludin, and ZO-1 penetrate Cx43 plaques. Example line scans show fluorescence in arbitrary units over the entire Cx43 plaque and perinexus region. Arrows indicate voids in the Cx43 plaque. **B)** Average line scans of relative fluorescence intensity for respective membrane proteins and Cx43 at the edge of Cx43 plaques. n ≥ 6. **C)** Respective Pearson correlation, or intermingling tendency, of membrane proteins with Cx43 within the gap junction plaque.

### Integration within Cx43 GJ plaques requires specific connexin domains

The overlapping distribution of Cx43 within gap junction plaques with Cx30, Ocln and the scaffolding protein ZO-1 raised the issue of whether the infiltration of the plaque was affected by structural rigidity of Cx43 within the plaque. To test this, we compared their normal distribution with that obtained in cells expressing highly fluid Cx43 plaques. The constructs we used deleted a large portion of the cytoplasmic tail of Cx43 or substituted three cysteine residues in this region with alanine; we previously showed that Cx43 is stabilized within the plaque by cysteine resides within its carboxyl terminus and that these mutations fluidize arrangement of channels with gap junction plaques made up of Cx43^6^.

ZO-1 associates with wild type Cx43 plaques (Figs 2A, 3A), but the association is variable, sometimes localizing at higher concentration near the perimeter of gap junction plaques. A similar, variable pattern of ZO-1 distribution is seen in plaques of fluid Cx43 (cysteine-substituted; BCx43cyslessCT) (Figure 3C). By contrast ZO-1 was not detected at plaques formed of truncated Cx43 (BCx43t258 in Figure 3B), although faint EGFP-ZO-1 signal was present outside of the Cx43 plaque area.

**Figure 3.**
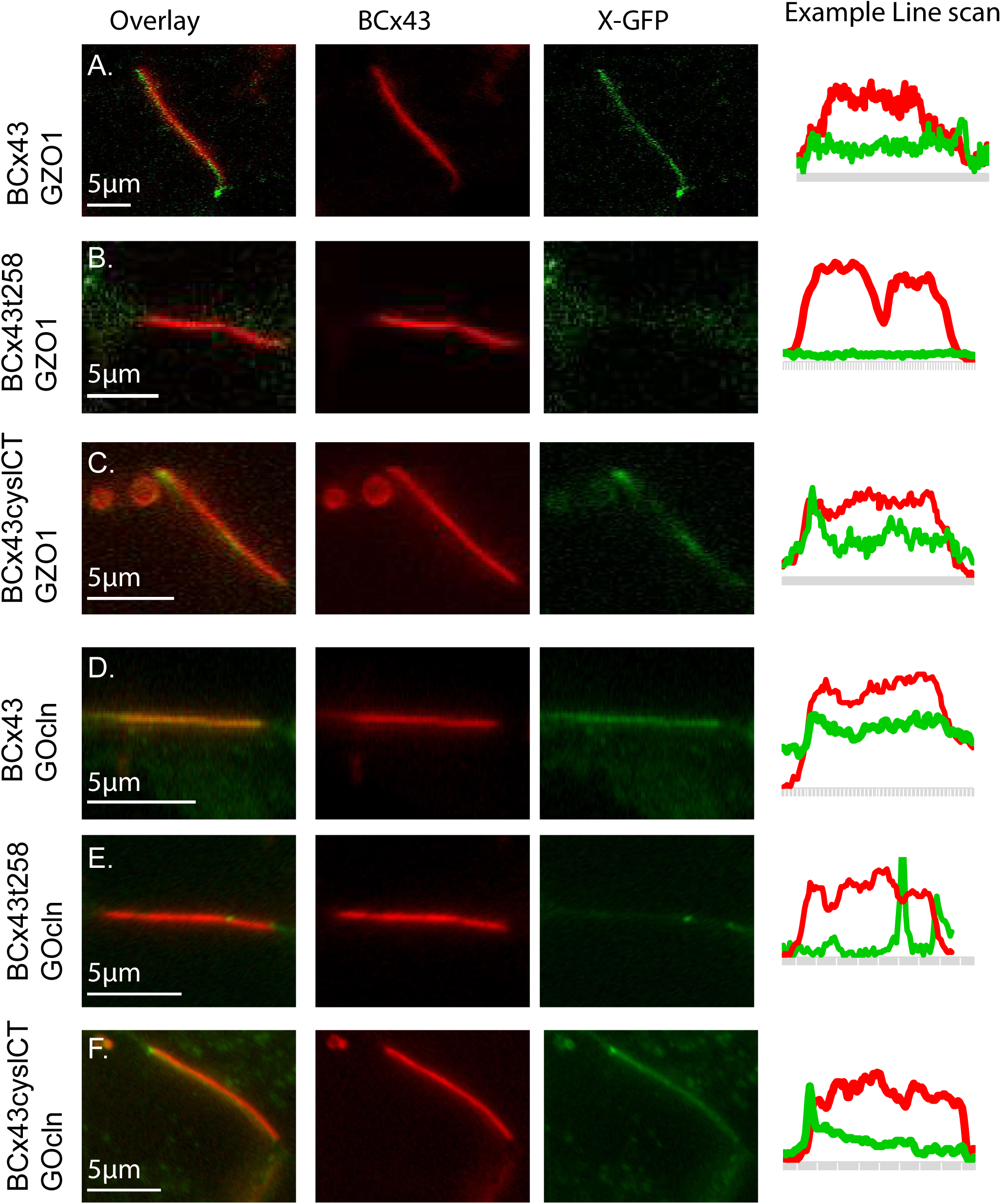
The attraction of some proteins to the gap junction plaque requires sequences in the carboxyl-terminus of Cx43. **A)** HeLa cells co-expressing EBFP2-Cx43 and EGFP-ZO1. Middle and right images show individual channels. The example line scan shows fluorescence in arbitrary units for the profile along the gap junction plaque and perinexus areas adjacent to the plaque edges. Fluorophore brightness and laser power varies greatly between the two channels and the intensity profiles are scaled for the line scan to allow spatial comparison. **B)** EBFP2-Cx43t258 (truncated at AA258, also known as K258stop) produces gap junction plaques but doAe.s not localize EGFP-ZO1 to the junction. **C)** EBFP2C-.Cx43cyslCT (C260A,_D_C. 271A, C298A) also produces gap junction plaques and EGFP-ZO1 localization to the plaque. The circular structures are endocytic vesicles (connexosomes) that are also sometimes observed in cells expressing the Cx43 form shown in “A” and “B”. **D)** EBFP2-Cx43 (full-length, wild-type rat Cx43 with blue tag, pseudocolored red for visibility) co-expressed with mEmerald-Occludin leads to localization of Occludin to the gap junction plaque. **E)** mEmerald-Occludin is not localized to the EBFP2-Cx43t258 gap junction plaque. **F)** EBFP2-Cx43cyslCT (with carboxyl-terminus intact) leads to enhanced localization of mEmerald to the gap junction plaque.

Interaction of Ocln with Cx43 also depended on the presence of the Cx43 carboxyl-terminus (Figure 3D-F). We found that EBFP2-Cx43 co-expression with mEmerald-Ocln (Figure 3D) generally led to strong concentration of Ocln specifically to the gap junction plaque, with some variability between cells and between plaques within the same cell. This attraction to the gap junction plaque was reversed (Ocln signal was less at the gap junction plaque than surrounding membrane) when truncated or carboxyl-terminus tagged Cx43 was expressed in conjunction with tagged Ocln (Figure 3E). Ocln association to the GJ plaque had the same requirements for the Cx43 carboxyl-terminus as ZO-1, suggesting that the PDZ binding site within the Cx43 cytoplasmic tail is required for colocalization of Ocln into the gap junction plaque.

Although there are Cx43-based gap junctions connecting astrocyte cellular processes throughout the parenchyma, Cx43 and AQP4 localization is concentrated to the perivascular astrocyte endfeet. This arrangement allowed us to examine distribution of the two proteins *in situ* using Stochastic Optical Reconstruction Microscopy (STORM) in fixed rat brain tissue sections and immunofluorescence staining (Figure 4). In this example image there is a striking lack of overlap between Cx43 and AQP4 staining around a transected brain blood vessel. These representative example data indicate that, at least in the case of AQP4, the findings of gap junction effects on protein localization identified in cell culture likely reflect the *in situ* condition.

**Figure 4.**
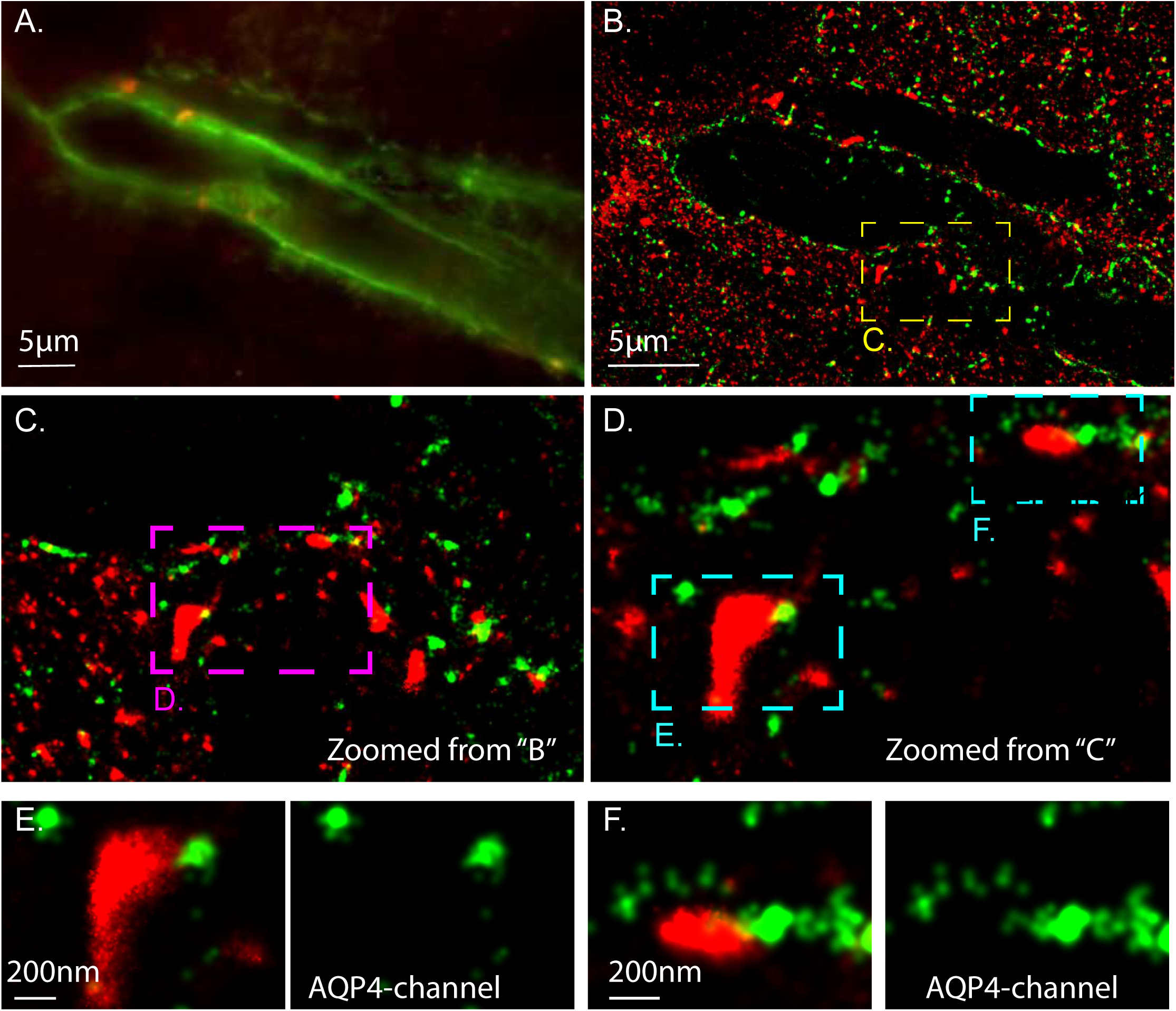
Non-overlapping Cx43 and AQP4 staining in rat astrocyte endfeet is revealed by two-color STochastic Reconstruction Optical Microscopy in immunostained cryosectioned tissue. STORM localization microscopy with two color channels (Alexa Flour 555 and Alexa Flour 647) in fixed rat brain tissue. A) Standard resolution wide-field microscopy. Aquaporin4 (green) is expressed in astrocyte endfeet around a blood vessel in the rat cortex. Connexin43 (red) forms gap junctions that connect astrocyte endfeet. B) Slightly zoomed in STORM image of the same brain blood vessel. More Cx43 molecules are visible outside of the astrocyte endfeet due to the ability to detect single Cx43 molecules with dSTORM and antibody labeling. C) Zoomed-in view from “B” on a region where astrocyte endfoot processes meet over the vessel. D) Further zoomed-in image of “C” to show large clusters of Cx43 signal in red are likely gap junction plaques. E and F) When zoomed-in beyond resolution of standard microscopy the lack of intermingling of the red Cx43 and green AQP4 signal is evident. AQP4 only (green) channel is shown to the panel to the right of the 2-channel overlay in “E” and “F”.

The fluorescent molecules examined here showed various patterns of distribution relative to the gap junction plaque and within the non-junctional membrane areas of the cells. We used Fluorescent Recovery After Photobleaching (FRAP) to determine their mobilities in non-junctional membranes and to compare their mobilities in stable vs fluid gap junction plaques. The fluorescent tag-labeled molecules each had differing levels of signal intensity within the non-junctional membrane due to several factors including fluorescent tag color, expression level of tagged proteins, and differential trafficking, cellular localization, and molecular clustering. Therefore, we present the FRAP data on plasma membrane over the same time-scale and photobleach area to allow rough comparison of mobility in non-junctional membrane with the caveat that numerous other parameters are likely not precisely matched between molecular species.

### Non-plaque Mobility

As shown in Figure 5, the rate of fluorescent recovery and percent recovery at 15 s post photobleach were markedly different between molecule types. Comparison of percent recovery at 15 s post bleach can be made to non-junctional msfGFP-Cx43 (likely existing as unpaired connexons, aka hemichannels, in a format of hexameric, 4-pass integral membrane proteins).

**Figure 5.**
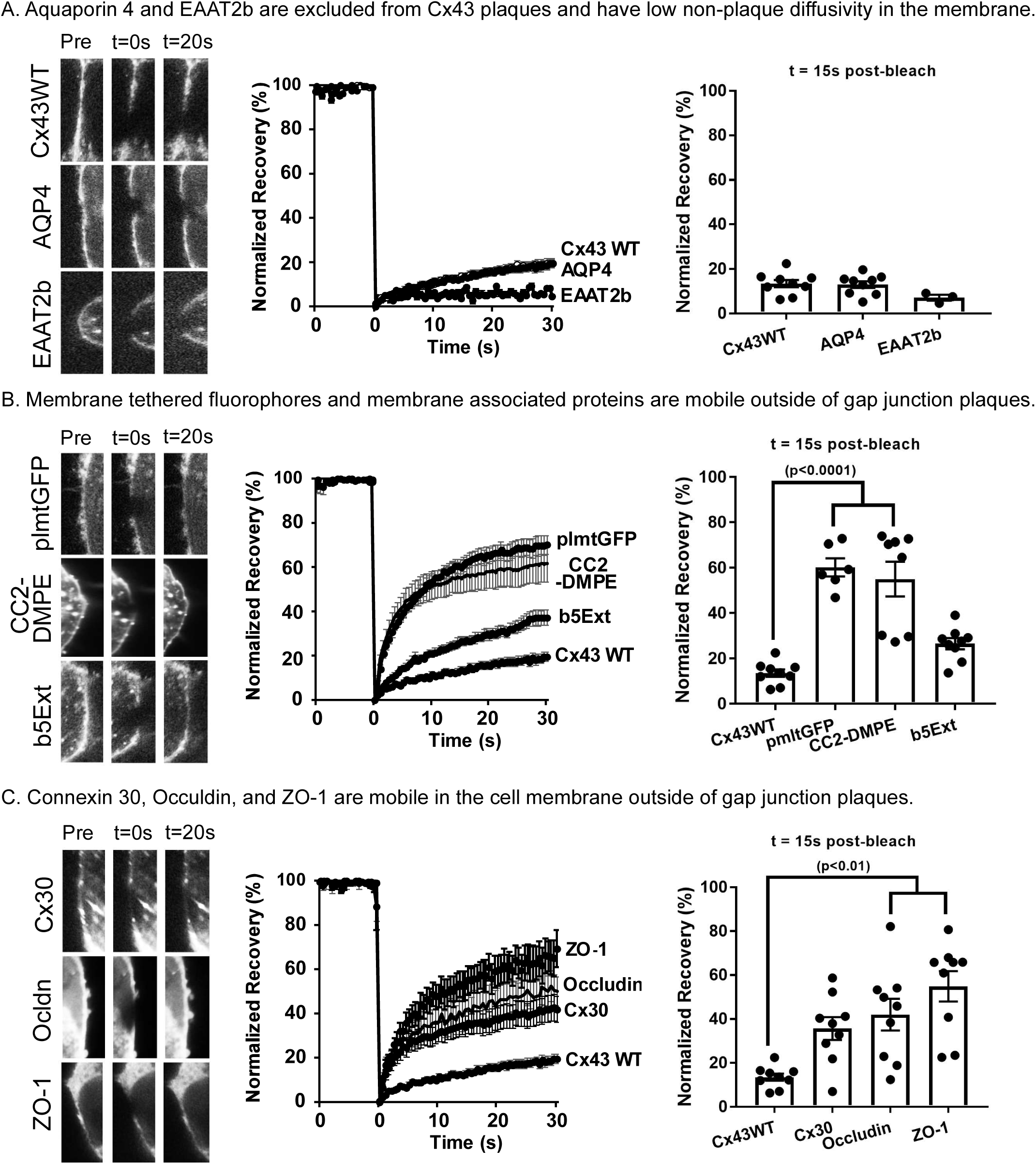
Nonplaque mobility of membrane embedded molecules. **A)** The plaque-excluded proteins aquaporin 4 and EAAT2b have low non-plaque membrane diffusivity. **B)** Membrane tethered fluorophores and membrane associate proteins are mobile outside of the gap junction plaque. **C)** Cx30, Occludin and ZO-1 are mobile in the cell membrane outside of the plaques. Each panel includes representative bleached non-plaque regions of interest, FRAP recovery curves, and bar chart of extent of recovery at 15 s post-bleach. Data shown as average ± SEM.

#### a) The plaque-excluded proteins AQP4 and EAAT2b have low non-plaque membrane diffusivity

Both AQP4 and the glutamate transporter EAAT2b were mostly excluded from gap junction plaques (Figure 2). FRAP experiments revealed that mobility of these proteins in non-junctional membrane was very low, and similar to that of wild type Cx43 outside of GJ plaques (Figure 5A), with neither of these proteins or unpaired Cx43 recovering more than about 20% within 30 sec after bleaching. Membrane effective diffusion coefficients for these molecules were 0.06±0.05 µm^2^/s for EAAT2b, 0.18±0.1 µm^2^/s for AQP4, and 0.24±0.06 µm^2^/s for unpaired Cx43 (Table 1, n=3).

**Table 1:** Diffusion coefficients calculated as defined in methods for each of the molecules studied in this report. Data presented as mean ± standard deviation. n ≥ 3. ^a^p<0.05 vs Cx43 wild-type (stable) plaque.

| | Diffusion Coefficient ($\mu\text{m}^2/\text{s}$) | | |
| --- | --- | --- | --- |
|  | Non-plaque | Stable Plaque | Mobile Plaque |
| Cx43 | $0.24 \pm 0.05^a$ | $0.03 \pm 0.01$ | $0.44 \pm 0.19^a$ |
| AQP4 | $0.18 \pm 0.10$ | n/a | n/a |
| EAAT2b | $0.06 \pm 0.05$ | n/a | n/a |
| CC2-DMPE | $4.38 \pm 1.05^a$ | $4.45 \pm 1.40^a$ | $3.48 \pm 0.68^a$ |
| plmtGFP | $4.84 \pm 2.58^a$ | $5.17 \pm 0.63^a$ | $4.54 \pm 2.90$ |
| b5Ext | $3.17 \pm 0.69^a$ | $3.02 \pm 0.52^a$ | $2.03 \pm 0.95$ |
| Cx30 | $1.43 \pm 0.35^a$ | $0.78 \pm 0.36$ | $1.85 \pm 1.35$ |
| Occludin | $2.11 \pm 0.55^a$ | $0.81 \pm 0.63$ | $2.51 \pm 0.92$ |
| ZO-1 | $1.49 \pm 0.31^a$ | $0.33 \pm 0.04^a$ | $0.44 \pm 0.62$ |

#### b) Membrane tethered fluorophores and membrane associated proteins are mobile outside of GJ plaques

The lipid CC2-DMPE and membrane tethered palmitoylated GFP located in non-junctional areas showed very rapid recovery from photobleach, with more than 50% recovery within 15 sec (54.9±20.3% for CC2-DMPE and 60.1±9.0% for plmtGFP: Fig 5B). By contrast, recovery was much slower for b5Ext, with only 26.4±6.8% recovery at 15 s post-bleach. Calculated effective diffusion coefficients were 4.84±2.58, 4.38±1.05 and 3.17±0.69 µm^2^/s for plmtGFP, CC2-DMPE and b4Ext, respectively (Table 1, n≥3).

#### c) Junctional proteins are mobile in the non-plaque region of the cell

Cx30, Ocln and ZO-1 co-mingle with Cx43 in gap junctions (Figure 2), and each was found to exhibit mobilities in the plasma membrane that were nearly as fast as those of the tagged phospholipid and the membrane embedded small molecules that we examined. Repopulation of the bleached area was greater than 35% complete within 15 s of each of these proteins; 35.7±14.7% for Cx30, 42.0±20.4% for Ocln, and 54.9±19.6% for ZO-1 (Figure 5C). Membrane effective diffusion coefficients were 1.43±0.35, 2.11±0.55 and 1.49±0.31 µm^2^/s for Cx30, Ocln and ZO-1 respectively (Table 1, n=3).

### Intraplaque Mobility

The colocalization experiments indicated a variety of overlap patterns between Cx43 and each of the different types of nexus components (Figure 1 and 2), as would be expected from binding to other proteins (membrane-embedded and cytoplasmic) and crowding within the highly ordered plaque. In order to investigate dynamics of these interactions we evaluated mobilities of those proteins in wild type and fluidzed Cx43 plaques using two color FRAP experiments.

#### a) Lipids and small integral membrane proteins are mobile within Cx43 GJ plaques

Due to differences in fluorescent signal density, photobleach region size, and photobleaching rate, comparisons of non-plaque and intraplaque mobility are not useful. However, the ability to modulate plaque stability by truncation or mutation allowed comparison of mobilities of the small lipophilic molecules in stable vs fluid plaque domains, revealing both similarities and distinctions (Fig. 6). The voltage sensitive membrane dye CCM2-DMPE exhibited very rapid recovery of about 50% of the bleached region within 15 sec, and this recovery was similar in stable (52.3±7.2%) and unstable (70.3±4.4%) GJ plaques (Fig 6A). Rates of recovery for CCM2-DMPE yielded effective diffusion coefficients of 4.45±1.4 and 3.48±0.68 µm^2^/s in stable and unstable plaques, which were not different from the non-junctional diffusion rate of 4.38±1.05 µm^2^/s (Table 1, n=3). Green Fluorescent Protein tethered to the membrane by palmitoylation also showed rapid recovery in both fluid and non-fluid plaques (Fig 6B). Recovery exceeded 60% within 15 sec (63.2±3.8% in non-fluid plaques and 71.6±4.2% in fluid plaques), and membrane effective diffusion coefficients were 5.17±0.63 and 4.54±2.9 µm^2^/s in non-fluid and fluid junctions, respectively, not different from rate in non-junctional membrane of 4.84±2.58 µm^2^/s (Table 1, n=3). The third lipophilic probe examined, b5Ext, also showed similarly rapid recovery of more than 40-50% within 15 sec (46.2±4.6% for WT and 57.7±6.6% for fluid plaques), with molecular diffusion coefficients of 3.02±0.52 and 2.03±0.95 µm^2^/s in WT and fluidized Cx43 plaques (Table 1, n=3). The extent of recovery was very similar to that in nonplaque domains for this molecule (3.17+0.69 µm^2^/s).

**Figure 6.**
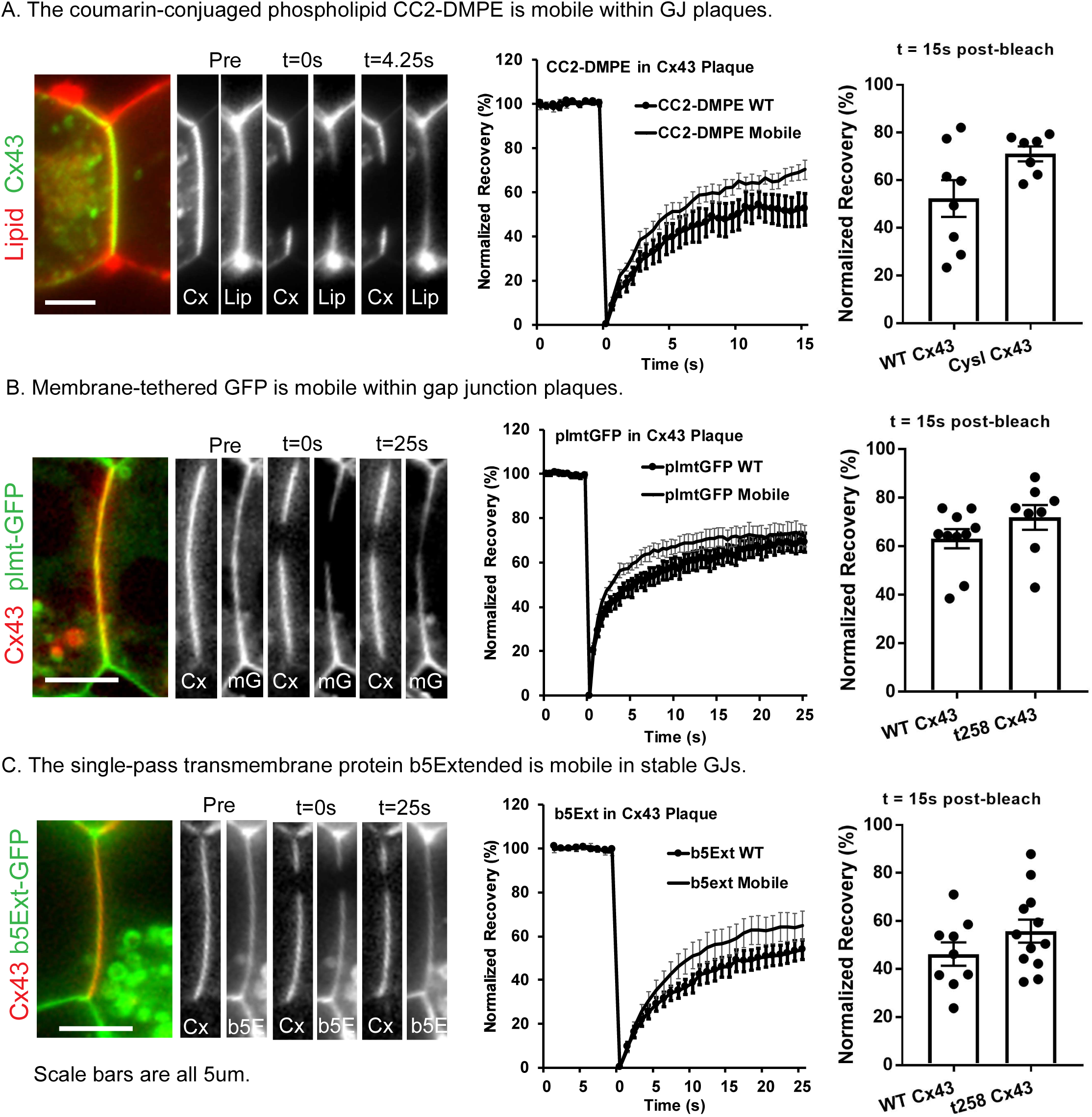
Lipids and small integral membrane proteins are mobile within both stable and fluid GJ plaques. **A)** The coumarin-conjugated phospholipid CC2-DMP is mobile within GJ plaques. **B)** Membrane-tethered GFP is mobile within GJ plaques. **C)** The single pass transmembrane protein b5Extended is mobile in stable GJs. Each panel includes representative bleached regions of interest, FRAP recovery curves, and bar chart of extent of recovery at 15 s post-bleach. Data shown as average ± SEM.

#### b) Behavior of junction-associated proteins in fluid an non-fluid GJ plaques

Both Cx30 and Ocln localize to the junctional membrane area occupied by gap junction plaques, ZO-1 is also located at GJ plaques (Figure 2), and we found that mobility of each of these proteins was relatively high in the non-junctional membrane (Fig 5C). Mobility of Cx30 within GJ plaques was higher and recovery was more complete in fluid than non-fluid (WT carboxyl-terminus) plaques. Recovery at 30 s post-bleach was 33.7±6.2% in WT plaques and 69.4±7.0% in fluidized plaques; diffusion coefficients were 0.78 ± 0.36 µm^2^/s in WT vs 1.85 ± 1.35 µm^2^/s in fluid plaques (Figure 7A).

**Figure 7.**
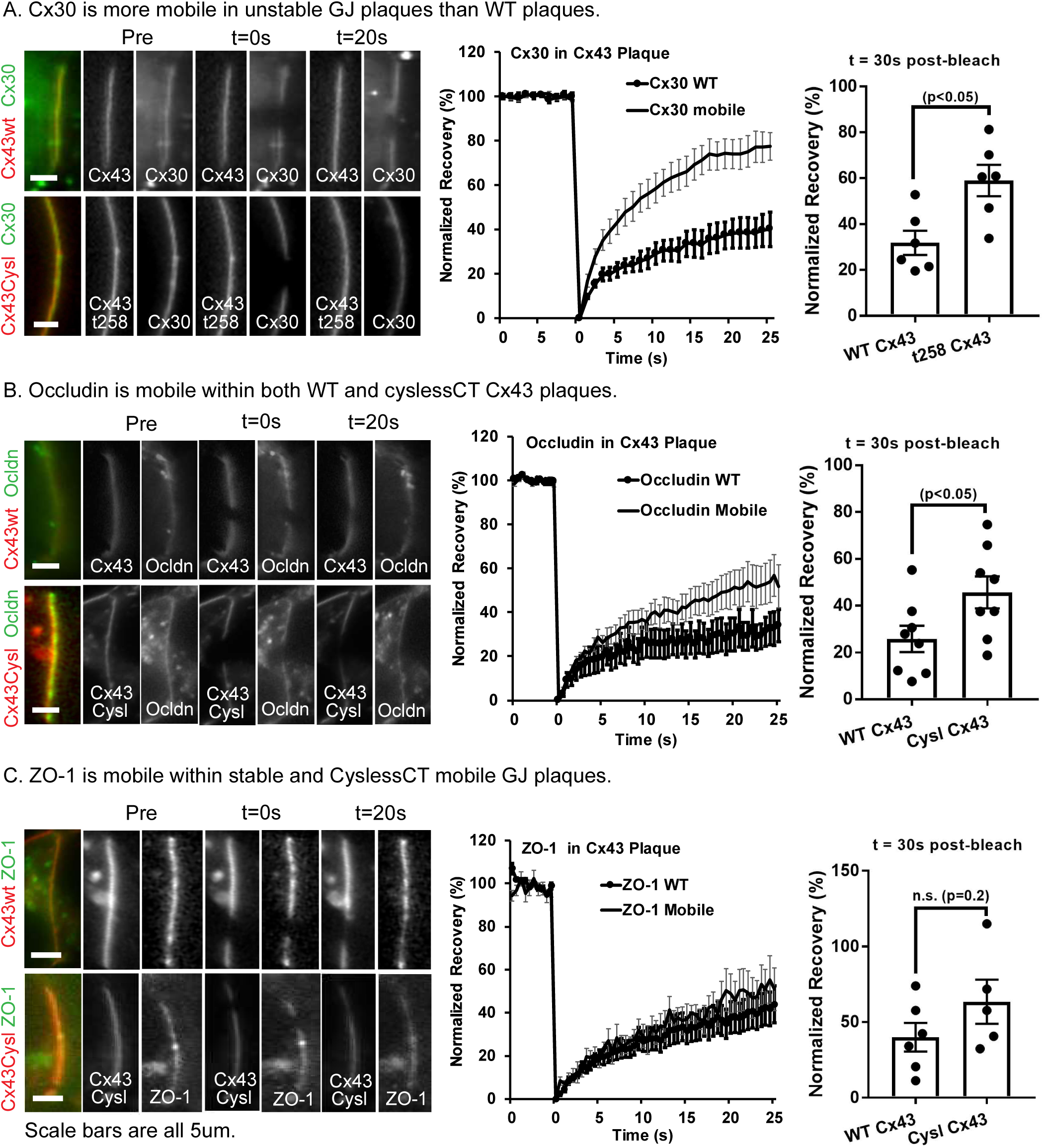
Junction-associated proteins are mobile in Cx43 GJ plaques and mobility is higher in fluid plaques. **A)** Cx30 is mobile within GJ plaques and is more mobile in unstable GJ plaques. Occludin is mobile within GJ plaques and is more mobile in unstable GJ plaques. **C)** ZO-1 is mobile within stable GJ plaques and does not localize to unstable GJ plaques. Each panel includes representative bleached regions of interest, FRAP recovery curves, and bar chart of extent of recovery at 15 s post-bleach. Data shown as average ± SEM.

We demonstrated that Ocln concentrates to Cx43 gap junction plaques but is excluded from plaques in which the Cx43 carboxyl terminus is deleted (Figure 3). FRAP experiments revealed that Ocln was much less mobile in normal Cx43 plaques than in fluid plaques, with recovery at 30 s post-bleach being 28.4±7.3% for WT plaques and 44.1±6.7% for cyslessCT (fluid) plaques. Effective diffusion coefficient in the non-fluid plaques was 0.81 ± 0.63 µm^2^/s, whereas mobility in fluid plaques (2.51 ± 0.92 µm^2^/s) was similar to that in non-junctional regions (2.11 ± 0.55 µm^2^/s, Figure 7B).

ZO-1 was found both at the edge of GJ plaques and also in a patchy distribution across the interior regions of wild type Cx43 gap junction plaques, but it was not localized to plaques made of truncated Cx43 (Figures 2 and 3). In photobleach experiments, ZO-1 recovered substantially less when in non-fluid junctions (31.7±6.1%) than in non-junctional membrane and effective diffusion coefficient was about 5 times slower (0.33 ± 0.04 µm^2^/s inside the plaque vs 1.49 ± 0.31 µm^2^/s outside). The diffusion coefficient of ZO-1 in cyslessCT (fluid) Cx43 plaques (0.44 ± 0.62 µm^2^/s was similar to that observed in non-fluid plaques with a trend toward increased mobility in the cysteine-mutant fluid plaques as shown in Figure 7C.

## Discussion

The highly-packed, two-dimensional array of connexons observed by freeze fracture and immunofluorescence of gap junction plaques does not reflect the many cytoplasmic and membrane proteins or lipophilic molecules that comprise the environment of the gap junction Nexus. Gap junction plaques made up of Cx43 with fluorescent protein tags have been observed bending, splitting, and joining with other gap junctions in live cells^14^. Additionally, density of connexins throughout the plaque may be nonuniform, with connexin-free discontinuities often observed to exist and move within stably arranged gap junctions^15^. Clearly, the apparent 2D crystalline array of intramembrane proteins belies the dynamic structure of the GJ plaque, where Nexus components bind and unbind to one another and diffuse within the plaque.

In this study we examined distribution of Nexus components relative to that of Cx43 in plaques in live cells using fluorescence recovery after photobleaching (FRAP) to determine diffusional mobilities of the molecules both within Cx43 plaques and in non-junctional domains. In addition, distribution and mobilities were compared in fluidized Cx43 plaques, where mutagenesis of cytoplasm-localized cysteine residues or truncation of the carboxyl terminus transforms the ordinarily rigid plaque into one which exhibited much more rapid and extensive FRAP recovery ^4,6^. Molecules examined included three small lipophilic markers, (a fluorescent phospholipid that has been used to detect voltage changes in membranes (CC2-DPME)^16^, GFP anchored to the membrane through palmitoylation, and b5Ext, a plasma membrane targeted fragment of cytochrome-b5 which has been used as a tool for studying membrane transport and endoplasmic reticulum^17,18^, two proteins forming pathways for exchange of molecules across the cell membrane (the water channel protein AQP4 and the glutamate transporter EAAT2b), and three components of cell junctions,(the gap junction protein Cx30, the tight junction protein Occludin (Ocln), and the scaffolding molecule zonula occludens (ZO-1), which binds to integral proteins of both tight and gap junctions).

Previous studies examined lipophilic dye movement through gap junction plaque-occupied membrane and found a high degree of mobility^19^. The lipophilic markers used in the present study were also highly mobile in non-junctional membrane and in both WT (stable) and cysless (fluid) Cx43 plaques. The lipophilic dye CC2-DMPE density varied only slightly across the plaque as was reported previously by others for the lipophilic DiI (1,1’-Dioctadecyl-3,3,3’,3’-Tetramethylindocarbocyanine Perchlorate)^19^. In contrast to CC2-DMPE, the other two membrane associated proteins we examined showed partial exclusion. We attribute the apparent exclusion as reflecting variation in lipid abundance between the general non-junctional membrane and the densely packed plaque^20^. With respect to mobility, the lipophilic molecules showed high non-junctional effective diffusion coefficients and mobility within the fluid plaque that were quite similar to that in non-fluid plaques.

Two transmembrane proteins, water channel AQP4 and glutamate transporter EAAT2b, were excluded from junctional plaques formed by both WT and cysless Cx43. Interestingly, AQP4 abundance consistently appeared higher in the region adjacent to the plaque than elsewhere (the “perinexus” ^21^) as shown in example line-scans and averaged line-scans in Figure 2 and in STORM experiments illustrated in Figure 4. Lateral mobility of these proteins in the membrane outside the plaque was about ten-fold lower than for other membrane proteins of similar size examined (Cx30 and Ocln). Previous studies have also reported low membrane mobility of AQP4 and EAAT2b and have speculated that stabilized AQP4 assembly into orthogonal particle arrays (OAPs) anchored by syntrophin may provide a mechanism for polarized water flux at astrocyte endfeet^22^, and that binding of EAAT2b to PDZ domain containing scaffolding molecules may optimize local glutamate uptake by astrocytes near synapses^23^.

The gap junction protein Cx30 and the tight junction-associated proteins Ocln and ZO-1 behaved differently than AQP4 and EAAT1b. All exhibited penetration within the junctional plaques, although abundance of Cx30 increased while ZO-1 was only minimally present in plaques formed by the truncation mutant. All displayed high mobility in regions outside the plaque, with diffusion coefficients on the same order of magnitude as those of the small lipophilic markers. Each had significantly decreased mobility within non-fluid plaques made up of Cx43 with wild-type carboxyl terminus. Mobility for Cx30 and Ocln was found to be higher in fluid than in non-fluid Cx43 plaques.

Ocln was the first ZO-1 binding partner identified, and the two proteins bind to one another to form a major component of tight junctions^24^. Ocln has previously been found to localize to gap junctions ^25,26^. We found that EBFP2-Cx43 or msfGFP-Cx43 co-expression with Ocln-mEmerald led to strong concentration of Ocln specifically to the gap junction plaque (Figure 2). This apparent attraction to the gap junction plaque was reversed (Ocln was partially excluded from GJ plaque) when truncated or a carboxyl-terminus tagged Cx43 was expressed with tagged Ocln (Figure 3), indicating that the PDZ binding site at the CT of Cx43 (the location where ZO-1 interacts)^27^ may be required for Cx43-induced localization of Ocln into GJ plaques. Cx43 has previously been shown to modulate tight junction formation ^28^ and this interaction with scaffold protein ZO-1 and Ocln may be one mechanism by which Cx43 promotes tight junctions, acting as a site for molecular aggregation and/or assembly.

In some conditions Cx43 GJ plaques are present in the basolateral membrane of epithelial cells where they are adjacent to (and sometimes intermingled with) tight and adherens junctions ^28-30^. We tested if gap junction plaque stability affected the localization and mobility of tight junction proteins. Cx43 binds ZO-1 through interaction between amino acids at the end of the carboxyl-terminus of Cx43 and the PDZ-2 binding domain of ZO-1 ^11,12,31,32^. Cx43 with a fluorescent tag appended to the carboxyl-terminus is unable to interact normally so we used our EBFP2-Cx43 construct with its free carboxyl-terminus to visualize the localization of Cx43 and ZO-1 in live cells ^11,33^. We found that while ZO-1 associates in the highest concentration at the edge of Cx43 gap junction plaques, as has been previously described^34^. ZO-1 was also reproducibly localized in a patchy distribution across the interior regions of wild type Cx43 gap junction plaques, but not localized to plaques made of truncated Cx43 (Figures 2 and 3). This could be, in part, an artifact of the overexpression of Cx43 and ZO-1 but indicates that ZO-1 binding is not strictly limited to the perimeter of the gap junctions. Notably, mEGFP-hZO-1 was found to be mobile at stably arranged gap junctions (Figure 7). Because ZO-1 is a cytosolic protein, photobleach recovery might occur from unbound unbleached protein in the vicinity of the bleach or from unbleached protein released from the prior location and rebound within the bleached region. We observed that recovery often proceeded from the edge of the bleach region toward the center of the bleach area as shown in Figure 5. This indicates that a large portion of the pool of unbleached mEGFP-hZO-1 that participated in photobleach recovery was from unbinding and re-localization of ZO-1 from bleach-adjacent parts of the GJ plaque. The preferential but incomplete localization of ZO-1 to the edge of gap junctions may be dependent on posttranslational modifications such as phosphorylation of Cx43 or could indicate competition with ZO-1 binding proteins^35,36^; actin and other junction proteins are likely candidates for such a factor promoting enhanced localization of ZO-1 to the edge of the plaque structure.

This distributed localization of ZO-1 allowed us to perform high-resolution FRAP experiments on ZO-1 localized to the gap junction nexus for the first time with the surprising result of relatively high mobility for ZO-1 (Effective Diffusion Coefficient 0.33±0.04 µ^2^/s, Table 1) on the intracellular surfaces of stably arranged Cx43 gap junctions. The amount and localization of ZO-1 and Ocln are dynamically altered and affect rate and degree of tight junction degradation and reformation, processes critical to diseases of barrier dysfunction at the interface between blood and brain, liver, kidney, and tumor (among others). The ability of Cx43 to concentrate or retain the tight junction protein Ocln at gap junctions has been observed previously^37^ but we now show that Ocln clustering into gap junction plaques depends on the carboxyl-terminus of Cx43 (Figure 2 and 3). The Cx43 gap junction Nexus may therefore form a platform for the assembly and retention of scaffolds and other junctional proteins. A previous study found that application of inflammatory cytokines to human astrocytes in culture modulated expression of claudins and tight junction proteins^38^. This raises the interesting question of whether Cx43 localization of ZO-1 and Ocln is modified by inflammatory cytokines as a potential alternative mechanism of tight-junction protein availability in astrocyte endfeet.

Although they are not necessary for the formation of gap junction plaques (channel clustering) cysteine residues within the Cx43 carboxyl-terminus act as tethers which hold clustered gap junctions in a stable arrangement. Connexin-free zones which appear as holes in gap junction plaque are generated when part of the gap junction is endocytosed^39^. The composition of these connexin-free zones is unknown, but they can migrate within an otherwise stable gap as described previously^40^. This movement of lipids and integral membrane proteins is perplexing in the context of a stably arranged gap junction but might be explained by weak and transient but extremely locally-abundant interactions as would be expected in the highly crowded and spatially ordered gap junction plaque. In the case of the CT of Cx43 gap junction channels, thirty-six cysteine residues (in total with paired- or “docked”-hexamers each containing 3 cytoplasmic cysteines) are positioned very close to cysteines on twelve immediately adjacent Cx43 gap junction channels. Additionally, the movement of the cysteines within the CT are restricted by the transmembrane domains and, likely, steric crowding by CT of adjacent Cx43 hexamers. Our results indicate that the anchoring behavior of the Cx43 CT restricts the mobility of other gap junction nexus components separately from the localization of these proteins to the gap junction plaque (which is dependent on a distinct sequence in the Cx43 CT downstream from the “Anchoring Domain” created by the three cysteine residues).

Cx43 and Cx30 gap junction plaques within the brain are localized to specialized astrocyte membrane extensions called endfeet where they cluster prominently around blood vessels and are also non-randomly localized to peri-synaptic astrocyte processes ^41-43^. Cx43 does not pair with Cx30 to form heterotypic junctions ^44^, although it can form GJ plaques with intermingled channels in some conditions and cell types ^4,41^ Cx43 and Cx30 are expressed in astrocytes where knockout studies indicate a complex but connexin-specific relationship to astrocyte- and by extension-brain function. Disruption of the gene for Cx43 alone or in combination with Cx30 had an opposite effect on synaptic signaling amplitude as Cx30 deletion, and the effect of Cx30 deletion was found to be channel-function independent^45^. Cx43 and Cx30 at astrocyte perivascular endfeet are required for normal blood-brain barrier strength or maintenance^46^. Along with extreme differences in the stability of the gap junction plaques that Cx30 forms, Cx30 also has a very different protein half-life, channel properties, and connexon assembly pathway within the cell compared with Cx43. It is possible that the effects Cx43 has on Cx30 arrangement within gap junctions could tune the parameters of half-life and channel open/closure depending on the tissue type (i.e. if the cells making up the tissue express only Cx30 or both Cx43 and Cx30).

Altogether, we report several new aspects of the gap junction Nexus supramolecular complex and that these new characteristics influence each other in surprising ways. Gap junction plaque arrangement stability lowers mobility of the transmembrane proteins we tested (Cx30 and Ocln) that intermingle into both stably and unstably arranged gap junctions. The cytoplasmic tight junction associated protein ZO-1 is known to interact with the CT and here we show that, surprisingly, fluorescent protein tagged ZO-1 is mobile when localized to stably arranged gap junction plaques. Lipid dyes were previously found to be mobile within stably arranged gap junction plaques ^19^. We extend these findings to show that that a synthetic dye-conjugated lipid, membrane tethered GFP, and a GFP-tagged single pass transmembrane protein are highly mobile within stable gap junction plaques. These results contribute substantially to our understanding of how the gap junction nexus is dynamically configured in live cells. Cx43 is co-expressed with Cx30 in astrocytes where the two proteins seem to have only partially overlapping roles in controlling intercellular communication and cell morphology. The role of cytoplasmic-localized cysteine residues in gap junction plaque stability, the effect of an antioxidant on gap junction plaque fluidity, and the interaction between Cx43 stability and other nexus components will be critical to further investigations to understand how gap junctions control cell/tissue morphology and physiology.

## Materials and Methods

### Plasmids and fluorescent probes

sfGFP-Cx43, sfGFP-Cx43t258, sfGFP-Cx30, Cx30-msfGFP, EBFP-Cx43, EBFP-Cx43t258 were described previously^4^. sfGFP-Cx43cyslCTwas described previously^6^.

The EGFP-ZO-1, Occludin-mEmerald and Occludin-mCherry expression plasmids were obtained from Addgene.com. pEGFP ZO1 was a gift from Alan Fanning (Addgene plasmid # 30313)^47^. pCAG-mGFP (palmitoylated GFP) was a gift from Connie Cepko (Addgene plasmid # 14757)^48^. The mCherry-Occludin-N-10, mEmerald-Occludin-N-10, and mCherry-Cx43-N-7 were gifts from Michael Davidson (Addgene plasmids # 55112, # 54212 and # 55023, respectively). EAAT2a-EGFP and EAAT2b-EGFP^49^ were gifts from Susan Amara, NIMH, Bethesda MD. The b5Extended-mGFP construct was a gift from Erik Snapp, Janelia Research Campus, Ashburn VA, and was originally described by Bulbarelli and colleagues^50^. The coumarin labeled phospholipid CC2-DMPE was a gift from Ted Bargiello, Albert Einstein College of Medicine, Bronx, NY.

### Cell culture and transfection

Neuro2a (N2A) and HeLa cells were maintained in DMEM medium (Glucose 4.5 g/L) supplemented with 10% FBS and 1% Penicillin/Streptomycin. For standard FRAP experiments N2A and HeLa cells were plated into 8-well imaging chambers (iBidi, cat no. 80826 or In Vitro Scientific, C8-1.5H-N) and each well was transfected with 0.5 μg of each plasmid to drive expression of Connexin-fluorescent protein fusions 24-72 h prior to imaging. Cells were transfected at ∼80% confluency. Optifect (Life Technologies) was used as the transfection regent according to manufacturer’s instructions adjusted for the surface area of the wells of the iBidi chambers (50 µl of Opti-MEM media and 3 µl of Optifect reagent per well). Opti-MEM media was replaced with the standard growth media for HeLa and N2A cells (DMEM with 10% fetal bovine serum and 1% Penicillin-Streptomycin 6-16 h after transfection. For experiments with co-expression of multiple proteins from separate plasmids (e.g. EBFP2-Cx43 + Cx30-msfGFP), the plasmids were mixed 1:1 prior to addition to the Opti-MEM-Optifect transfection mixture. Cells were incubated in standard growth media for at least 4 h prior to imaging. For non-plaque FRAP experiments transfection was formed using TransIT-LT1 (Mirus, LLC, Madison, WI) per manufacturer’s protocol. 2 µg of DNA, 100 µL of Opti-MEM, and 5 µL per 3.5 mm dish were mixed and incubated at room temperature for 30 min and added dropwise to HeLa cell cultures. Experiments were performed at 24-72 h post-transfection.

### Confocal microscopy and line-scans

2D confocal micrographs were obtained with the Zeiss LSM 510 Live with Duo module with 63x NA=1.4 oil immersion objective. Images were 512×1024 or 1024×1024 pixels. An imaging plane was selected with a view of the gap junction plaque orthogonal to the cell membrane, such that plaques appeared as linear elements. Single frame images were taken and analyzed for molecular distribution as quantified by fluorescence intensity. Line scans over entire Cx43 plaques, starting approximately 1-2 µm outside of the plaque, were obtained using ImageJ (NIH), by tracing plaque contours with the line tool and measuring intensity profile. Pearson correlations were calculated for each Cx43-plaque associated molecule pair^51^.

### Two-color Stochastic Optical Reconstruction Microscopy (STORM)

Brain of adult rats were extracted immediately after the rats died and placed in 4% Paraformaldehyde in PBS for 48 hours at 4°C. The brains were transferred to 30% sucrose for 96 hours at 4°C then frozen into Optimal Cryosectioning Tissue gel and cryosectioned to 15 μm in the coronal orientation through the forebrain in the area including the hippocampus. Sections were placed free-floating in PBS and immediately put through the immunostaining procedure. Immunostaining was performed by first blocking in permeabilizing blocking buffer and background suppression buffer from Biotium (TrueBlack® IF Background Suppressor System 1:100 dilution in PBS, Cat no. #23012-T). Primary antibodies were polyclonal Goat anti-AQP4 (Cat. No. sc-9888, Santa Cruz Biotechnology) and rabbit anti-Cx43 (Cat. No. C6219, Sigma Aldrich). Primary antibodies were diluted to 1:500 in blocking buffer, sections were incubated with agitation for 3 hours at room temperature followed by 24 hours at 4°C, followed by 3 hours at room temperature. Sections were washed in PBS once then transferred to secondary antibody at 1:500 dilution in blocking buffer for 3 hours at room temperature with agitation. Sections were washed in PBS for 5 minutes at room temperature with agitation 3 times, then post-fixed in 2% PFA for 10 minutes, washed once in PBS then stored until imaging (1-2 days).

Imaging was performed on the Nanoimager S (Oxford NanoImaging, ONI, Oxford, UK). Sections were mounted onto coverslips in PBS then a drop of BCubed STORM buffer (ONI) was placed onto the section followed by overlay of a #1.5 coverslip and the coverslip sealed with clear fingernail polish. The sections were imaged using the 100X 1.41NA Olympus objective on the Nanoimager at 37°C at 100% laser power onto the dual-color channel sCMOS camera. Localizations image display was performed in the NimOS v1.3 software. The signal from the Alexa Fluor 555 and 647 was acquired with 100 percent laser power simultaneously for 20,000 frames. Localizations were shown by the “Precision” method which makes displayed spot size and blur depend on localization precision.

### FRAP

2D time-lapse imaging was conducted as described previously ^4^, and ^52^. Cells were maintained at 37°C on the stage of a Zeiss LSM 510 Live with Duo module and imaged with a 63X NA=1.4 oil immersion objective. The detector consists of dual 512 pixel linear arrays of CCD camera-type pixels. Gap junctions aligned in a nearly perpendicular plane with respect to the growth substrate were used for 2D time-lapse FRAP. A 5 pixel (1 μm wide) stripe bleach region was set to bleach a horizontal stripe across each gap junction plaque. Bleach settings were 100% laser transmission at a scan speed of 5 with 3 bleach iterations. Lower bleach laser power and single bleach iterations were tested to generate greatly reduced photobleaching. No recovery of Cx43 into the bleach region (no detectable rearrangement of GJ channels) was observed in FRAP experiments with the lowest degree of bleaching. This indicates that photobleach induced oxidation is not the cause of Cx43 stability.

### Time-lapse FRAP

Experiments in which only GFP was bleached use the same procedure as normal 2D time-lapse FRAP with 1 s acquisition intervals instead of 0.5s intervals but with sequential excitation-detection scanning with the GFP (green) channel and then the EBFB2 (blue channel, shown as red in all figures for visibility) as described ^52^. FRAP recovery data for the blue and green channels were extracted separately as described below. Laser power and detector gain sometimes needed to be adjusted within samples of the same group.

### FRAP data analysis

Average fluorescence within the bleach region, for the entire GJ plaque to be bleached (Fluorescence pool available for recovery, Fp), and a portion of the background in a location with no GFP expression were outlined to generate 3 Regions Of Interest (ROI). Recovery curves were transformed to correct for loss of signal due to bleach and for acquisition-bleach of the total pool of fluorescent protein and normalized to 100% pre-bleach and 0% for the initial post-bleach time-point to normalize for incomplete bleaching within the bleach ROI as previously described ^53^. A correction factor (cf) was calculated by dividing the average of the 10 fluorescence pool readings preceding the bleach (initial fluorescence of the fluorescence pool; fpFo) by the fluorescent pool ROI readings at each time point (Fp), (fpFo/Fp). The bleach ROI reading (bF) for each time point was divided by the bleach region baseline-initial fluorescence Fo (bFo), (bF/bFo) and the resulting fraction of initial fluorescence was then multiplied by the correction factor. The resulting corrected fractional fluorescence was then multiplied by 100% to calculate “normalized recovery (%).” With omission of background subtraction, transforming to complete bleach baseline, and averaging to generate initial fluorescence pool values, the calculations for the recovery curve values were as follows:

FRAP%=(fpFo/Fp)*(bF/bFo)*100%.

The normalized data points at 15 or 30 s after the bleach time-point were used in comparison of percent recovery at 15 or 30 s. Effective diffusion coefficient was estimated using the ImageJ/FIJI FRAP plugin “simFRAP” which fits the data to a simulated diffusion ^54-56^. In brief, simFRAP loads image data into a 2D diffusion simulation with user-defined ROIs: a bleach region, bleached cell region (recovery pool), and reference cell region. We followed the supplied protocol with modification of ROI selection to adjust for the shape of gap junction plaques. We used the bleached region of the plaque as the bleached ROI, but used 2 μm of the plaque on either side of the bleach region as recovery pool ROI, and a 2 μm section of the plaque distal from the bleach region (or a separate gap junction plaque) as the reference cell ROI. Simulations were run for 10000 iterations. Diffusion coefficient was fitted by the equation D_cal_=L^2^/4 τ_*i*_ where D_cal_ is the calculated diffusion coefficient L is pixel size in µm and τ_*i*_ is the duration of one iteration in seconds.

### Statistics

For comparison of 2 groups we used a student’s t-test. When comparing 3 or more groups we used a one-way AVONA with pair-wise post-hoc Tukey Tests. Significance between groups was defined as p < 0.05. Statistical analysis was performed using GraphPad Prism 7.03 for Windows (GraphPad Software, La Jolla California USA).

## Abbreviations

AQP4: aquaporin 4;
*OAP*: orthogonal array of particles formed by AQP4;
GJ: Gap Junction;
Cx: connexin;
Cx30: connexin 30;
Cx43: connexin 43;
EAAT2b: excitatory amino acid transporter 2b;
CC2-DMPE: coumarin phospholipid;
plmtGFP: palmitoylated GFP;
b5Ext: b5Extended;
ZO-1: zona occludens 1;
Ocln: occludin;
EBFP2: enhanced blue fluorescent protein 2;
FRAP: fluorescence recovery after photobleaching;
GFP: green fluorescent protein;
HEPES: 4-(2-hydroxyethyl)-1-piperazineethanesulfonic acid;
msfGFP: monomerized superfolder green fluorescent protein;
ROI: region of interest;
RT: room temperature;
sfGFP: non-monomerized superfolder green fluorescent protein;
GFP-Cx43t*XXX*: rat Cx43 tagged with GFP on the amino-terminus truncated at the indicated amino acid (i.e. GFP-Cx43t258 is truncated by mutagenesis of lysine 258 to a stop codon);
Cx43cyslCT: rat Cx43 with cysteine residues 260, 271, and 298 mutated to alanine;
TJ: tight junction;

## Acknowledgments

Super-resolution imaging was performed with support of the NYIT Imaging Center. The NYIT Center for Biomedical Innovation provided RFS with computational tools for image processing and analysis

